# The transcriptional mechanism behind *Mimosa pudica* leaf folding in response to mechanical disturbance

**DOI:** 10.1101/2025.07.17.665295

**Authors:** Matteo Buti, Alice Checcucci, Chiara Vergata, Luciana Renna, Susanna Pollastri, Francesco Loreto, Stefano Mancuso, Federico Martinelli

## Abstract

*Mimosa pudica* is a plant known for its ability to fold leaves in response to mechanical disturbances, which serves as a visible phenotypic stress marker. Leaf folding occurs with a response timing and intensity that varies depending on the stimulus. This adaptive behavior may function as a defense mechanism, helping plant resist herbivores and environmental stressors. However, the molecular and genetic mechanisms that are involved in leaf folding are still not totally understood. In this study, the gene regulatory networks underlying *M. pudica* leaf closure following single and multiple mechanical disturbances (whole pot drops) were investigated. Chlorophyll fluorescence was measured as fast phenotypic indicator of transient or permanent photochemical damage, and transcriptional responses were measured to identify the key genes regulating phenotypic changes after single or multiple drops. A progressive reduction of the quantum yield of photosystem 2 revealed a lower electron transport rate in leaves subjected to one or more drops, which may indicate the onset of energy shortage, perhaps caused by low ATP availability, limiting both leaf movement and photosynthesis. The transcriptomic profiles revealed larger differences when plants were subjected to multiple drops than to a single drop, with respect to unstressed controls. Interestingly, following a single drop, the majority of up-regulated genes were associated with the flavonoid biosynthetic pathway. After multiple drops, however, genes associated with biotic and abiotic stress resistance pathways were predominantly up-regulated. These results provide a basis for developing a gene regulatory network model of stress-induced movements in *M. pudica* leaves, which may help design sustainable strategies of plant stress defense.

**Main Conclusions:** Repeated stress in *Mimosa pudica* reduces photosystem efficiency, alters gene expression, shifting from flavonoid biosynthesis to stress resistance pathways, offering insights for sustainable plant stress defense strategies.

## Introduction

To adapt to their sessile nature, plants have evolved growth responses that enable them to cope with the frequent, repetitive, and rapid changes in their environment (Hagihara and Toyota 2020). Generally, the mechanism that enables organisms to respond quickly to external stimuli after achieving a physiological state that prepares them to cope with stress is known as an adaptive preparation process (Aranega-Bou et al. 2014 ; Harris et al. 2023). Several studies evidenced that plants are able of mnemonic functions regulating specific physiological and morphological responses, which are determined by genetic or epigenetic mechanisms (Kinoshita & Jacobsen 2012; He, Y., & Li, Z., 2018). Exploring the molecular and genetic dimensions of this alert state, including the initial perception of stimuli, the swift response to these stimuli, and the subsequent memory of past stress experience (Gagliano et al. 2014), is particularly intriguing.

Exposure to mechanical stimulation has long been known to influence plant growth and development, shaping both biomechanical properties of plants and their overall dimensions, according to the thigmomorphogenetic process (Chehab et al. 2009). A range of phytohormones and signaling molecules, such as intracellular calcium, reactive oxygen species, ethylene, jasmonates abscisic acid, brassinosteroids, auxin and nitric oxide, are known to be involved in touch responses. Following touch stimulation, numerous genes are activated, encoding proteins that participate in several cellular processes, including, cell wall modifications calcium sensing and defense mechanisms (Chehab et al. 2009). In particular, several genes and transcription factors (TF), referred to as touch-induced (TCH) genes, have been investigated for their role in perception and transduction of mechanical stimuli in various plant species (Braam and Davis 1990; Botella et al. 1996; Monshausen and Haswell 2013). Among these, Piezo genes encode mechanosensitive ion channels that play a crucial role in several physiological processes, including perception of mechanical disturbance and the consequent regulation of calcium ion flow (Kaur et al. 2023).

Numerous genes linked to responses to biotic and abiotic stressors may also be involved in mechanical stress responses. During signals’ transmission, intracellular calcium levels fluctuate, activating several calcium-dependent enzymes such as calmodulin and protein kinases. These enzymes subsequently trigger downstream signaling pathways that regulate numerous cellular processes (Lee and Seo 2021). Genes such as *NPR1* (Nonexpressor of Pathogenesis-Related Genes 1) (Dong 2004; Lai et al. 2018; Zavaliev and Dong 2024), and transcription factors from WRKY (Meraj et al. 2020, Chen et al. 2012), AP2/ERF (Mizoi et al. 2012; Wang et al. 2023) and DREB2 families, can also be involved in the maintenance of cellular homeostasis after both biotic and/or abiotic stresses and regulate stress responses as well as general plant memory mechanisms (Wang et al. 2016).

Additionally, genes involved in secondary metabolite production, such as those in phenylpropanoid pathway and flavonoid biosynthesis, are described as important for their antioxidant properties in stress response mechanisms (Falcone Ferreyra et al. 2012; Sharma et al. 2019).

Finally, DNA methylation and histone modification genes have been found to play a role in plant mnemonic processes, influencing transcriptional regulation of genes and the modification of chromatin structure (Li et al. 2007; Zhang et al. 2018; Liu and He 2020).

Two studies from the last decade explored the transcriptomic profiles of *Arabidopsis thaliana* (Lee et al. 2005) and *Populus nigra* (Fluch et al. 2008) following touch-stimulations, providing further details on the molecular mechanisms underlying their mechanical response. In these studies, the up- and down-regulated genes identified after a single time-point stimulation were mostly related to defense mechanisms, such as those involved in oxygen species (ROS) production, enzymatic ROS scavenging and in the phenylpropanoid pathway. Other studies investigated the kinetics of the transcriptional profile after one stimulation, although this cannot explain the response to multiple stimuli (Kimbrough et al. 2004). Two other researches demonstrated the existence of a desensitization process to mechanical stimuli involving transcriptional regulation (Martin et al. 2010; Pomiès et al. 2017). These studies showed that, following a second stress event, the group of genes that altered their expression differed from those that responded to a single mechanical stimulus, displaying a unique expression pattern, and influencing various biological processes. Indeed, short-term transcriptomic responses quickly activated abiotic stress pathways and plant defense mechanisms, such as ethylene and jasmonic acid signaling, as well as photosynthesis regulation, while later responses impacted genes related to cell wall formation and wood development.

*Mimosa pudica* is a plant known for its capability of folding leaves in response to stresses like local wounding, strong temperature variations, and mechanical stimulation (Bakshi et al. 2023). Leaf folding following contact disturbance has been identified as a defensive response to predators. Indeed, it seems sustained longer under high-light conditions compared to light-limited conditions (Jensen et al. 2011). An intriguing feature of *M. pudica* leaf closure behavior is that a touch stimulus can trigger a signal that spreads to nearby leaves, causing even those that were not touched to close (Malone 1994). However, the mechanisms of signals perception and transduction triggering such a rapid response remain unknown.

Recently, Hagihara et al. (2022) started to investigate the genetic basis of *M. pudica* leaf movement, along with the genetic and molecular characterization of the ion channels involved in the signaling system controlling leaf movements (Lee et al. 2005; Tran et al. 2021). Moreover, epigenetic regulation and the role of transcription factors have been recently explored in primed plants developing immunity or stress tolerance (Harris et al. 2023). Currently, the gene expression patterns associated with the leaf multi-folding response of *M. pudica* and the underlying biochemical mechanisms remain largely unexplored. However, these aspects could be of significant interest to get a deeper insight into how plants respond to mechanical stress.

In this study, the gene regulatory networks underlying *M. pudica* leaf closure in response to single and multiple mechanical disturbances were investigated. Potted plants were subjected to single or multiple downfalls that stimulated the rapid but temporary folding of leaves along the stem, followed by a return to the original unfolded conditions with a variable recovery time. Photosynthetic and transcriptional responses in plants exposed to one or multiple downfalls were compared to those of undisturbed plants to identify the key factors in gene expression regulation related to acclimation and memory of perceived mechanical disturbances.

## Materials and Methods

### Experimental set-up and design

Twenty-four plants at identical developmental stages (10 cm tall, 5 leaves brunches), 18 months old, were purchased from the local specialized nursery Vivaio Calicanto in Florence, and grown for a week for acclimatation in a growth chamber with a temperature of 23 °C, a 16/8 hours light/dark regime and 60% humidity. Three different treatments were applied to eight plants each. ‘Control’ plants were left non-stimulated, with their leaves wide open. ‘Stimulated’ plants were subjected to a single drop from a height of 15 cm, as described by Gagliano et al. (2014). ‘Overstimulated’ plants were subjected to repetitive drops from the same height, following the protocol by Gagliano et al. (2014), with cycles of leaf closing and reopening until the leaves no longer responded to the stimulus (adapted leaves).

Plants subjected to the three different treatments were either measured for chlorophyll fluorescence parameters or sampled for transcriptomic analysis. Samples were collected immediately (seconds) after the treatments in the case of a single stimulation (within seconds) and following all repetitive drops in the case of multiple stimulations (10 minutes from the start of the treatment). In this case the leaves of three plants for each treatment were cut at the hypocotyls base and instantly frozen in liquid nitrogen.

### Chlorophyll fluorescence

The Imaging Pam M-series fluorimeter (Heinz Walz) was used to measure chlorophyll fluorescence parameters in leaves. After the treatment, ‘control’, ‘stimulated’ and ‘overstimulated’ plants were first adapted to darkness for 20 min, then individually placed inside the fluorimeter chamber and photosystem II (PSII) maximum quantum yield in leaves was measured as the ratio between variable fluorescence and maximal fluorescence (Fv/Fm). Plants were then exposed to actinic light (110 μmol photons m^-2^ s^-1^) and, after 10 min, the PSII quantum yield in illuminated leaves (Φ PSII) was also measured according to Genty et al. (1989). For each treatment, measurements were performed on five biological replicates.

### RNA isolation and sequencing

Total RNA isolation was performed using approximately 50 mg of the immediately after stimulation frozen material ground with a pestle and mortar following the Norgen Plant/Fungi Total RNA Purification Kit protocol with slight modifications in the cell lysis phase. Briefly, Proteinase K was added to the Lysis Buffer C and β-mercaptoethanol mixture, and pulverized samples were heated at 56 °C for 10 min under bead beating condition. For the final elutions, the solutions were brought to 60 μL using RNase-free water. Maximum 10 μg of extracted RNA was treated with DNAase I (NEB) before quality check. RNA integrity was determined for each sample using the Agilent 2100 Bioanalyzer (RNA 6000 nano kit - Biorad Inc).

Sequencing libraries were obtained following the Stranded mRNA Library Prep procedure (Illumina) with an exclusive unique dual index combination (3 libraries for each of the three experimental conditions). Qubit™ 4 Fluorometer (dsDNA High Sensitivity Kit - Invitrogen) was used to estimate the concentration of the libraries, while quality assessment was performed by Agilent 2100 Bioanalyzer (DNA HS kit - Biorad Inc). Novaseq 6000 Reagent Kit (2x100 + 10 + 10 cycles) was used to perform sequencing, with all samples processed in a single flow cell.

### De novo assembly of the Mimosa pudica transcriptome

The RNA-Seq obtained from the ‘control’, ‘stimulated’ and ‘overstimulated’ plants were used to generate a *de novo* transcriptome assembly of *Mimosa pudica*. After the conversion of RNA-Seq data to fastq files with Illumina bcl2fastq v2.20, RNA-Seq raw reads quality was assessed with FastQC v0.11.5 (Andrews 2010), and adapters sequences and low-quality reads were removed with Trimmomatic v0.39 (Bolger et al. 2014) using these settings: ILLUMINACLIP:adapt.fa:2:30:10 HEADCROP:1 LEADING:3 TRAILING:3 SLIDINGWINDOW:4:18 MINLEN:40.

The RNA reads of all the libraries were used for a *de novo* assembling with Trinity v2.13.2 (Grabherr et al. 2011), and assembled transcripts were clustered using CD-hit v4.8.1 (Fu et al. 2012) to produce a set of ‘non-redundant’ transcripts. Statistics of non-clustered and clustered *de novo* assembled transcriptome were obtained with ‘n50’ Perl script from the SeqFu suite (Telatin et al. 2021). The final transcriptome completeness was assessed with BUSCO v5.3.2 (Simão et al. 2015) using the ‘viridiplantae_odb10’ lineage.

### Transcripts level estimation and differential expression analyses

Bowtie2 v2.4.4 (Langmead and Salzberg 2012) with default parameters was used to map back the reads of the sequenced RNA libraries were to the final *Mimosa* transcriptome, while the expression quantification of each transcript was carried out with the Salmon v1.4 (Patro et al. 2017) tools ‘quant’ and ‘quantmerge’.

Raw counts data elaborations and differential expression (DE) analyses were performed with Bioconductor EdgeR v3.38.1 (Robinson et al. 2009). This tool was used to filter out the not active transcripts (a transcript was considered ‘active’ if reads per million mapping was >1 in two or more libraries), normalize the RNA libraries according to their depth, visualize the multi-dimensional scaling (MDS) plot for RNA libraries normalized counts, and do the differential expression analyses. A transcript was considered as differentially expressed in a pairwise comparison if its false discovery rate (FDR) was lower than 0.05 and its log_2_ Fold Change (LFC) was lower than −2 or higher than 2. Venn diagram representations of differentially expressed transcripts (DETs) were visualized with Venny 2.1.0 (Oliveros 2015) and InteractiVenn (Heberle et al. 2015).

### Functional annotation and enrichment analyses

Functional annotation of DETs were performed on the Galaxy platform (Afgan et al. 2018) using the BLAST+ blastx-fast algorithm (Camacho et al. 2009). Searches were conducted against the NCBI nr, SwissProt, RefSeq, TrEMBL, UniRef100 Viridiplantae and Araport11 protein databases, applying an e-value cutoff of 10^-3^. To perform enrichment analyses, Gene Ontology (GO) terms were assigned to the entire assembled transcriptome of *M. pudica* using EggNOG (Cantalapiedra et al. 2021) with ‘Viridiplantae’ taxonomic scope. Additionally, KEGG orthologs were mapped using the (KEGG

Automatic Annotation Server) (Moriya et al. 2007) with BBH assignment method, using Arabidopsis and available Fabaceae species as reference organisms.

GO enrichment analyses was conducted in Cytoscape 3.10.1 (Shannon et al. 2003) using the BinGO 3.0.5 plug-in (Maere et al. 2005), with a hypergeometric test, FDR correction, a significance level of 0.05, and ‘GOSlim_Plants’ ontology. Enriched KEGG classes across the three pairwise comparisons were identified using clusterProfiler v4.6.2 (Wu et al. 2021), via a custom R script. KEGG classes were considered enriched in a comparison if the adjusted p-value was below 0.05, and the results were visualized using ggplot2 v3.5.1 (Wickham 2016).

## Results

### Response of Mimosa pudica plants to dropping

#### Leaves movement

Once subjected to dropping as described by Gagliano et al. (2014), ‘stimulated’ plants performed a folding movement of their leaflet within 30 s, soon after also actuating a dropping movement of the petiole (directly attached to the stem). The plant’s entire response to the dropping was usually completed within one min. ‘Overstimulated’ plants did not actuate any of the movements made by ‘stimulated’ plants.

#### Chlorophyll fluorescence measurements

To evaluate the photosynthetic performances of stressed plants, the quantum yield of PSII in leaves of ‘control’, ‘stimulated’ and ‘overstimulated’ plants *Mimosa* plants was measured by chlorophyll fluorescence. The maximal quantum yield of PSII in dark-adapted leaves was similar in all the samples (Fig. 1 A, B, C, D). Whereas, in illuminated plants, the PSII quantum yield was significantly higher in ‘control’ than in ‘stimulated’ and ‘overstimulated’ plants (Fig. 1, E, F, G, H).

**Figure 1.**
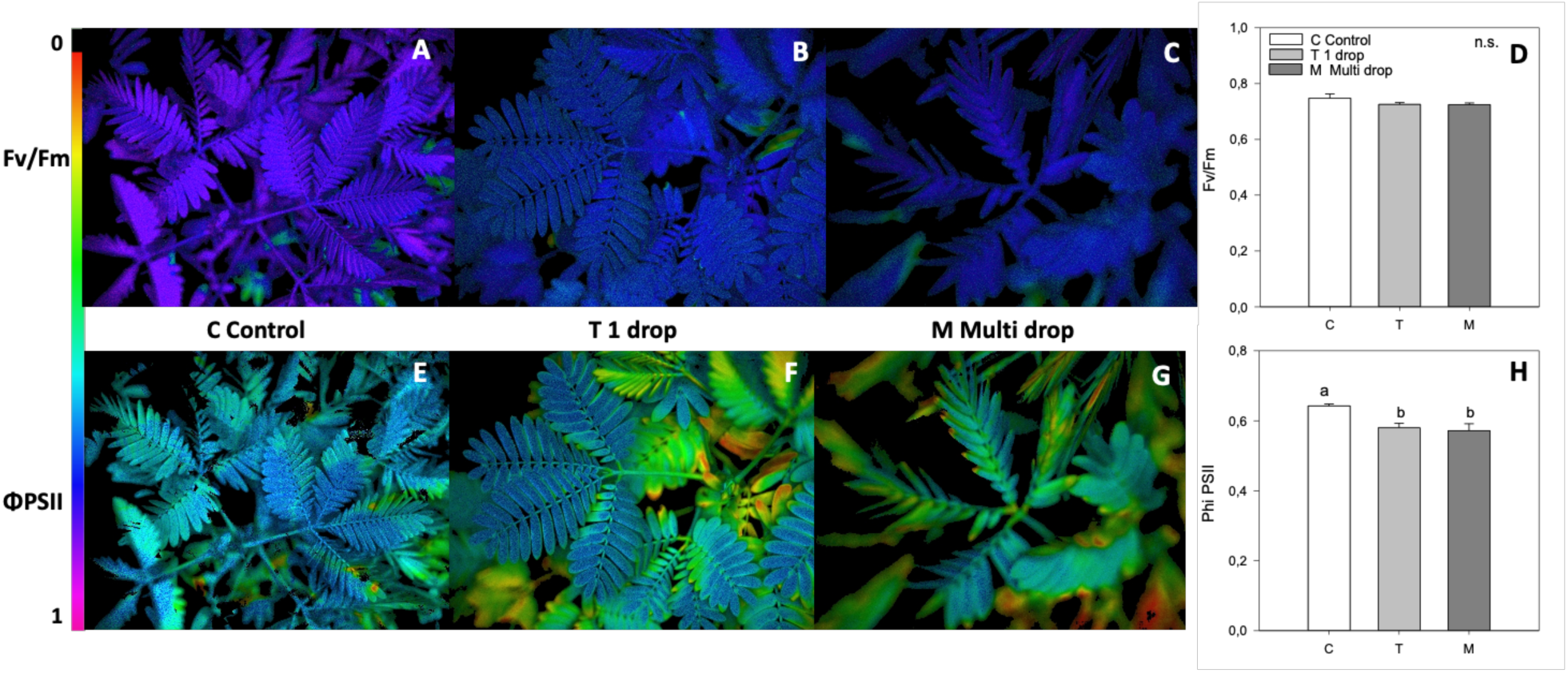
Representative chlorophyll fluorescence images of PSII maximum quantum yield (F_V_/F_M_) and PSII quantum yield (Φ PSII) in Mimosa pudica plants, in control conditions (C; A,E), after one drop (T; B,F) and after multidrop (M; C,G). Light adapted plants were subjected to no drop (C), 1 drop (T) or multi drop (M) treatment and then after 20 minutes in the dark, F_V_/F_M_ measurements were performed. Then plants were exposed to actinic light (110 PAR) and after 10 minutes Φ PSII was measured. The colour bar panels (D,H) showed means (*n* = 5) ± SE of FV/Fm and PhiPSII in control conditions (C, white bars), after 1 drop (T, light grey bars), and after multidrop (M, grey bars). A one-way ANOVA followed by Tukey’s test was performed to define statistically significant differences among means (*P* < 0.05). Means not sharing the same letters are statistically significantly different (n.s. not significant). The colour bar on the left of the panel shows the ranges of chlorophyll fluorescence.

#### RNA sequencing and transcriptome assembling

RNA libraries were sequenced, and the number of resulting paired reads was 120.2 million, ranging from 11.3 to 20.5 million for each library (Table 1). Due to the low sequencing quality of the C3 library, one ‘control’ sample was excluded from the downstream bioinformatic analyses. The RNA raw reads, once their quality was assessed, were deposited with the E-MTAB-14230 accession number in the ArrayExpress database. The percentage of ‘survived’ reads after the elimination of low-quality reads and adapters sequences ranged from 89% to 94% across the libraries (Table 1).

**Table 1.** RNA-seq data analysis. For each library, the number of raw reads, number and percentage of filtered reads, and number and percentage of reads mapped to the *de novo* assembled *M. pudica* transcriptome were reported.

| *Sample properties* |  |  | *Filtering* |  | *Mapping* |
| --- | --- | --- | --- | --- | --- |
| Treatment | Sample ID | Input read pairs | N° pairs survived | % pairs survived | % aligned pairs |
| ‘control’ | C1 | 13,703,838 | 12,701,613 | 92.69% | 90.90% |
|  | C2 | 20,467,660 | 18,807,858 | 91.89% | 91.97% |
| ‘stimulated’ | T1 | 17,913,579 | 16,543,531 | 92.35% | 93.68% |
|  | T2 | 10,287,641 | 9,447,847 | 91.84% | 93.54% |
|  | T3 | 19,158,762 | 17,455,579 | 91.11% | 93.89% |
| ‘overstimulated’ | M1 | 14,984,276 | 13,839,471 | 92.36% | 96.71% |
|  | M2 | 11,329,360 | 10,510,197 | 92.77% | 96.32% |
|  | M3 | 12,342,792 | 11,346,338 | 91.93% | 96.73% |

Filtered reads of the RNA libraries were *de novo* assembled, yielding 120,942 transcripts with an N50 of 1,736 covering 128,242,001 nucleotides. The assembled transcripts were then clustered to reduce redundancy, resulting in a final transcriptome of 93,662 transcripts covering 88,897,694 nucleotides. A BUSCO assessment of this transcriptome showed a high completeness score of 96.4%.

#### Analysis of transcriptional changes after mechanical disturbances

The RNA libraries reads mapping to the *Mimosa* transcriptome assembly ranged from 90 to 97% (Table 1). The mapping of reads to each transcript in the individual RNA libraries was then quantified. After filtering out the ‘not active’ genes (43,487 out of the 93,662 assembled transcripts were classified as ‘active’), normalization factors were assessed based on libraries size, and normalized reads counts were calculated. The MDS plot demonstrated high reproducibility, with biological replicates closely clustered and clearly distinct (Fig. S1).

DE analysis was performed for three pairwise comparisons: plants dropped one time *vs*. control plants (‘stimulated’ *vs.* ‘control’), plants dropped multiple times *vs*. control plants (‘overstimulated’ *vs.* ‘control’) and plants dropped multiple times *vs*. plants dropped one time (‘overstimulated’ *vs.* ‘stimulated’). The DETs for the three pairwise comparisons, along with their associated analysis results, were provided in Table S1. A summary of DET counts is presented in Table 2.

**Table 2.**
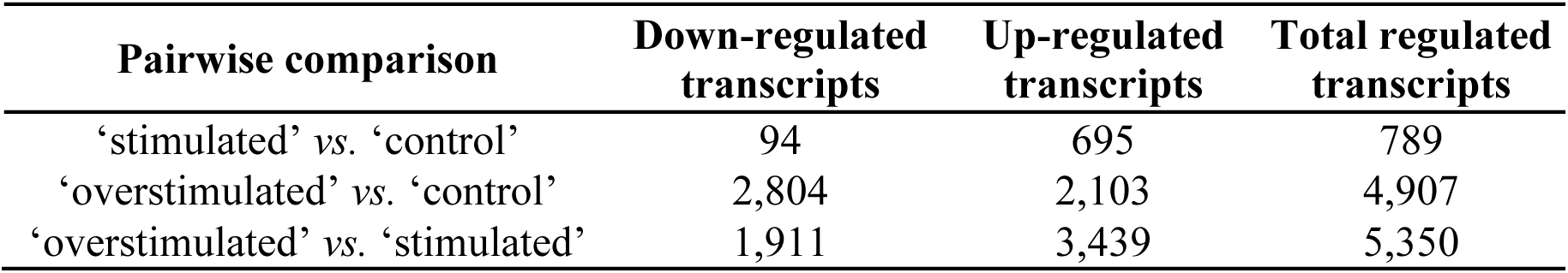
DE analysis results summary. For the three pairwise comparisons, the numbers of down-regulated, up-regulated and total regulated transcript were reported.

| Pairwise comparison | Down-regulated transcripts | Up-regulated transcripts | Total regulated transcripts |
| --- | --- | --- | --- |
| ‘stimulated’ vs. ‘control’ | 94 | 695 | 789 |
| ‘overstimulated’ vs. ‘control’ | 2,804 | 2,103 | 4,907 |
| ‘overstimulated’ vs. ‘stimulated’ | 1,911 | 3,439 | 5,350 |

A clustering heatmap of all the DETs was generated, revealing distinct transcript groups with varying expression patterns (Fig. 2A). In particular, this clusterization evidenced the similarities between the results of the comparisons involving plants dropped multiple times (‘overstimulated’ *vs.* ‘control’ and ‘overstimulated’ vs. ‘stimulated’). Single dropping shapes the expression level of 789 transcripts, among which 94 were down-regulated and 695 were up-regulated; multiple droppings induced a differential transcriptional activity in 4,907 transcripts compared to control plants, among which 2,804 were down-regulated and 2,103 were up-regulated; furthermore, multiple droppings induced a differential transcriptional activity in 5,350 transcripts compared to single dropped plants, among which 3,439 were down-regulated and 1,911 were up-regulated (Table 2). According to these numbers, single dropping induced transcriptomic changes in a smaller number of genes compared to multiple droppings, predominantly up-regulating them.

**Figure 2.**
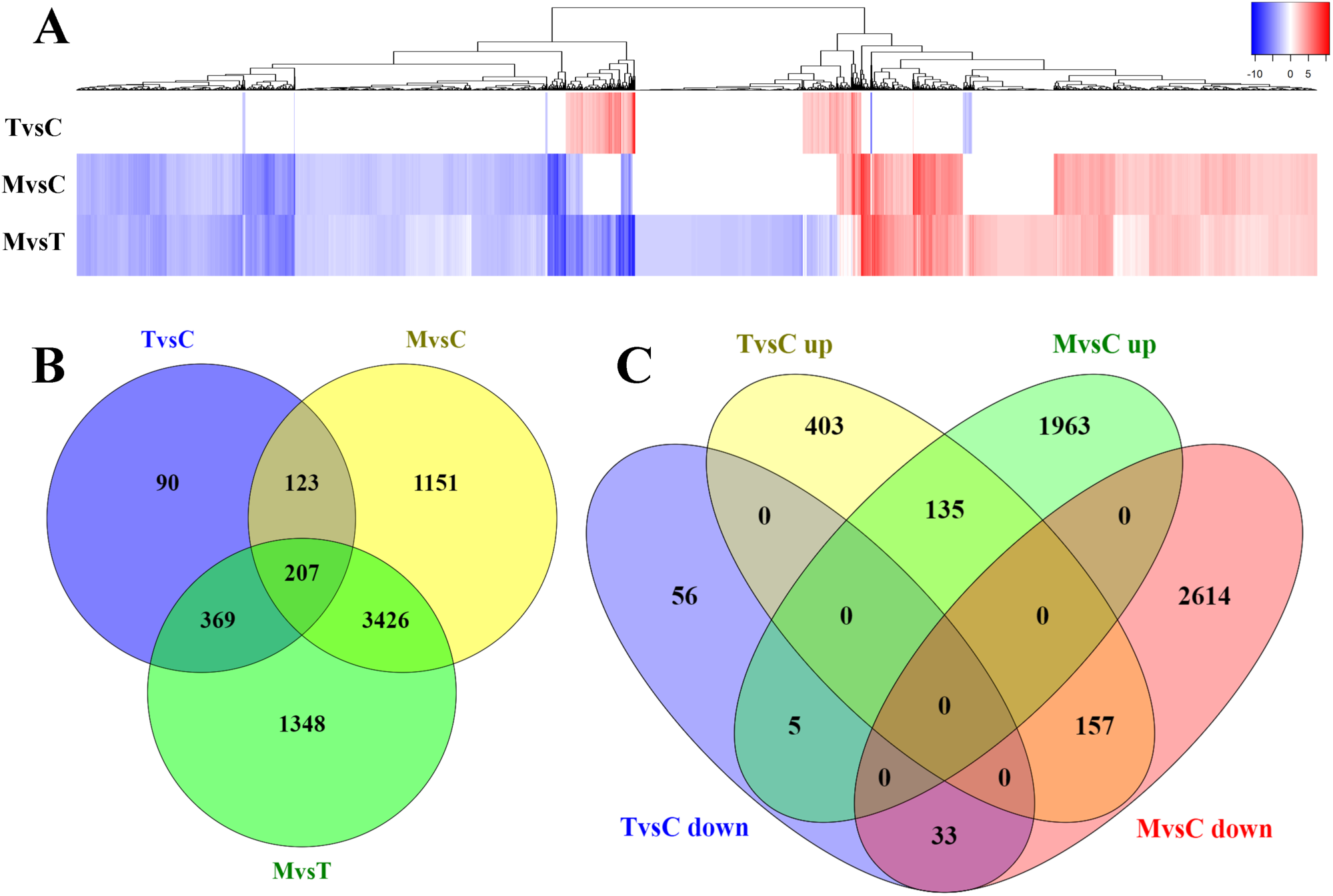
DE analyses results. A. Heatmap reporting the logFC of each DET for the three pairwise comparisons. Color key for logFC value was displayed; B. Venn diagram of DETs of the three comparisons ‘stimulated’ vs. ‘control’ (TvsC), ‘overstimulated’ vs. ‘control’ (MvsC) and ‘overstimulated’ vs. ‘stimulated’ (MvsT); C. Venn diagram of up- and down-regulated transcripts for the comparisons ‘stimulated’ vs. ‘control’ (TvsC) and ‘overstimulated’ vs. ‘control’ (MvsC).

Interestingly, the number of DETs is higher in the comparison between multi-dropping and single-dropping than in the other comparisons, even surpassing the number of DETs between multi-dropping and control plants. Furthermore, while the number of up-regulated DETs was greater than the number of down-regulated ones after a single dropping, the transcriptomic changes induced after multi-dropping consistently shifted towards the down-regulation in both ‘overstimulated’ *vs.* ‘control’ and ‘overstimulated’ *vs.* ‘stimulated’ comparisons. Surprisingly, this pattern is more pronounced in the multi-dropping *vs.* single-dropping comparison (Table 2), revealing a significantly larger number of expression differences than in the single-dropping *vs.* control plants comparison.

Indeed, as shown in Fig. 2B, the number of DETs shared between the comparisons involving multi-dropping are nearly ten times higher than those shared between the single-dropping and control plant comparisons (3,426 versus 369, respectively). Moreover, according to the Venn diagram of the comparisons ‘stimulated’ *vs.* ‘control’, and ‘overstimulated’ *vs.* ‘control’, (Fig. 2C), most of the differentially regulated transcripts are specific for each pairwise comparison, as evidenced also in 6-way Venn diagram in Fig. S2, where DETs are unique only in the comparisons involving ‘overstimulated’ samples. However, 135 and 33 transcripts that are up- or down-regulated in both comparisons were identified, respectively. Among these, transcripts with contrasting expression pattern in the two pairwise comparisons were found: 157 were up-regulated in ‘stimulated’ *vs.* ‘control’ and down-regulated in ‘overstimulated’ *vs.* ‘control’, and 5 were down-regulated in ‘stimulated’ *vs.* ‘control’ and up-regulated in ‘overstimulated’ *vs.* ‘control’.

Transcripts that resulted as differentially expressed in at least one of the three pairwise comparisons were functionally annotated. Of the 6,714 DETs, a total of 5,638 (84.0%), 5,631 (83.9%), 4,721 (70.3%), 5,634 (83.9%), and 5,362 (79.9%) returned at least one hit after the BLASTp analysis with the NCBI nr, RefSeq, SwissProt, TrEMBL and Araport11 databases as subjects, respectively. The detailed functional annotation results were presented in Table S2.

#### GO- and KEGG-enrichment analyses results

Enrichment analysis facilitates the classification of gene functions and the identification of over-represented biological processes and pathways, offering a comprehensive understanding of the underlying biological mechanisms. This enhances the interpretability and relevance of our transcriptomic data. Consequently, Gene Ontology (GO) and Kyoto Encyclopedia of Genes and Genomes (KEGG) terms were assigned to the entire *de novo* assembled *M. pudica* transcriptome to perform the required gene set enrichment analyses. Among the *M. pudica* assembled transcripts, GO terms were assigned to 14,833, while KEGG orthologs were assigned to 10,311 (Table S3). The number of enriched GO categories was 7, 14 and 16 for ‘stimulated’ *vs.* ‘control’, ‘overstimulated’ *vs.* ‘control’ and ‘overstimulated’ *vs.* ‘stimulated’ pairwise comparisons, respectively (Table 3).

**Table 3.**
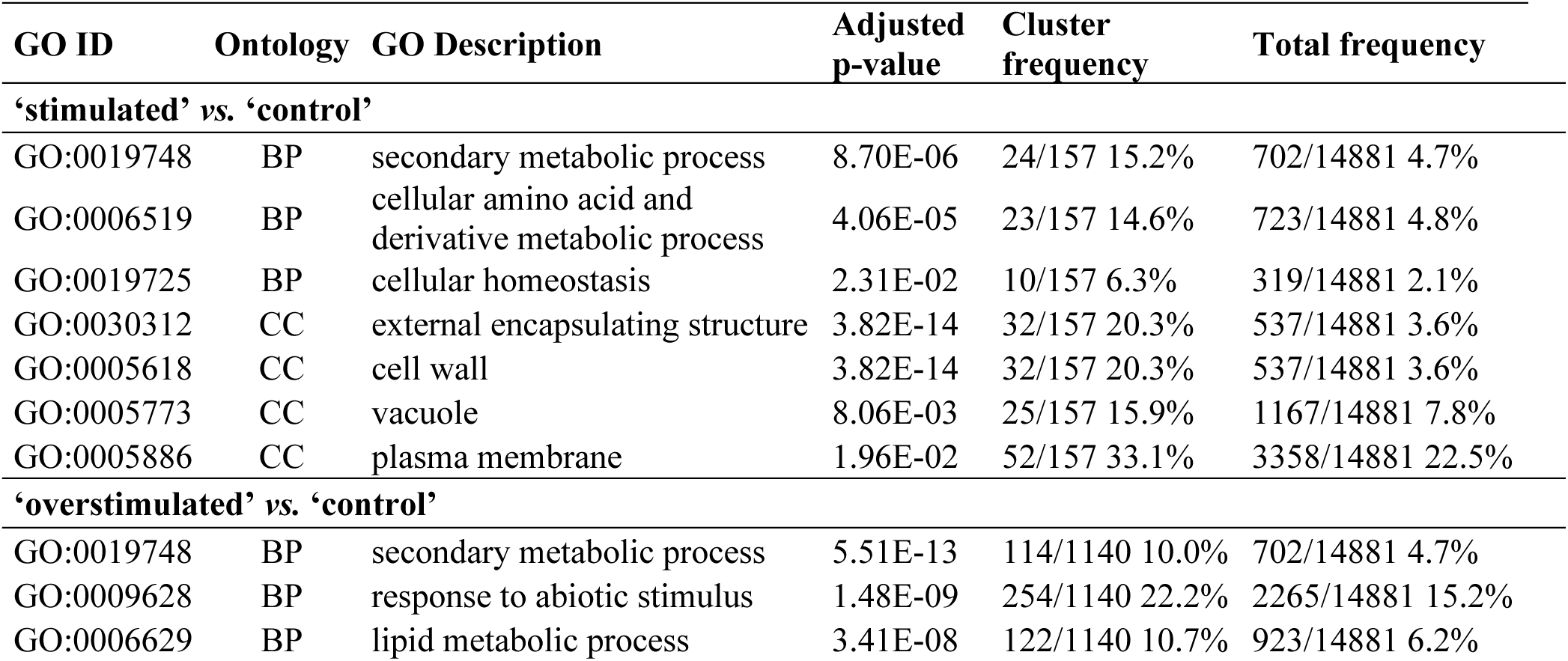
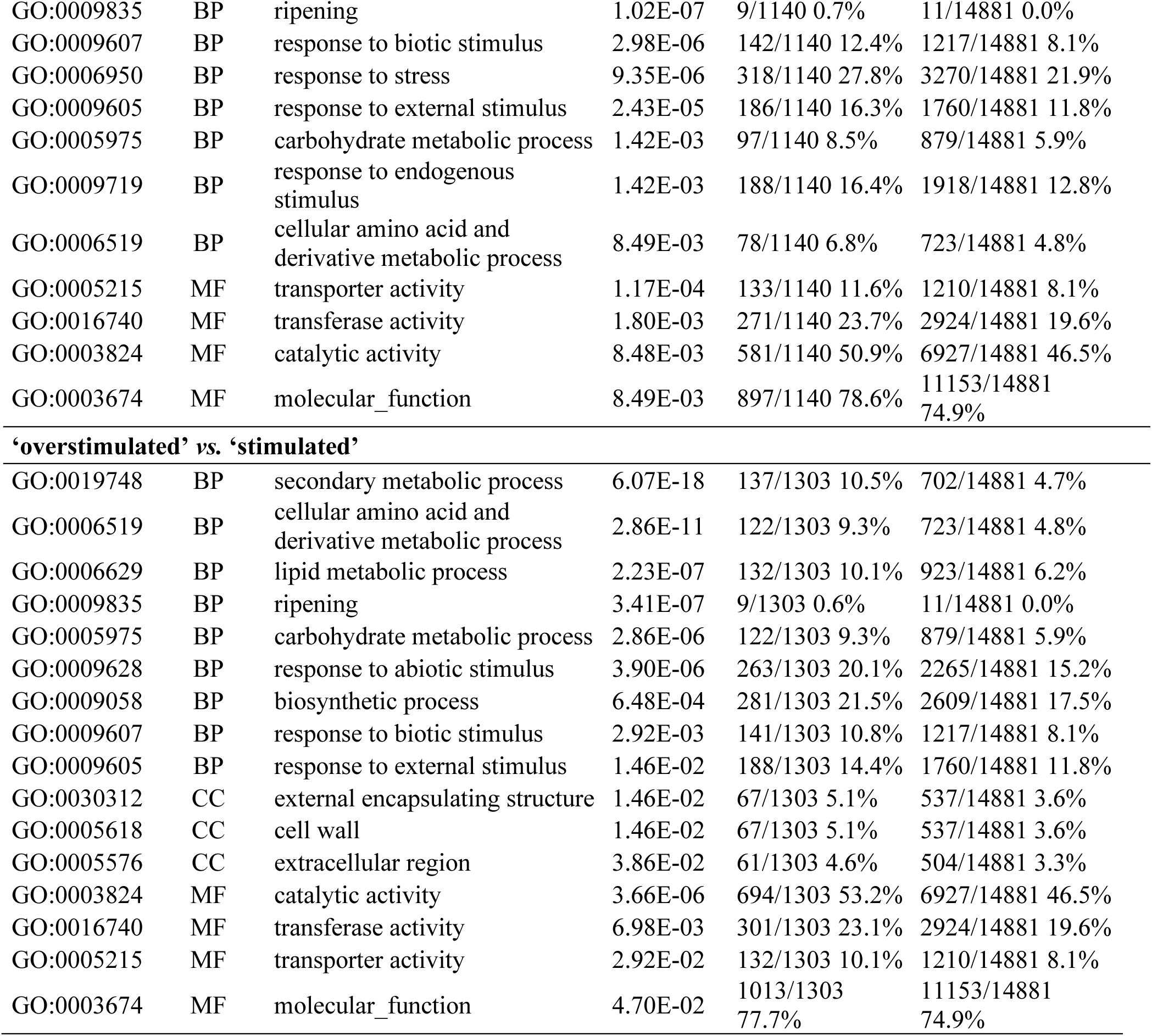
Enriched GO categories for DETs obtained from ‘stimulated’ *vs.* ‘control’, ‘overstimulated’ *vs.* ‘control’, and ‘overstimulated’ *vs.* ‘stimulated’ pairwise comparisons. Each record reports the GO term ID, its ontology (biological process (BP), molecular function (MF), cellular component (CC)), description, adjusted p-value, frequency of the GO term in the analyzed DETs (ratio and percentage), and frequency in the entire dataset (ratio and percentage).

| GO ID | Ontology | GO Description | Adjusted p-value | Cluster frequency | Total frequency |
| --- | --- | --- | --- | --- | --- |
| **‘stimulated’ vs. ‘control’** |  |  |  |  |  |
| GO:0019748 | BP | secondary metabolic process | 8.70E-06 | 24/157 15.2% | 702/14881 4.7% |
| GO:0006519 | BP | cellular amino acid and derivative metabolic process | 4.06E-05 | 23/157 14.6% | 723/14881 4.8% |
| GO:0019725 | BP | cellular homeostasis | 2.31E-02 | 10/157 6.3% | 319/14881 2.1% |
| GO:0030312 | CC | external encapsulating structure | 3.82E-14 | 32/157 20.3% | 537/14881 3.6% |
| GO:0005618 | CC | cell wall | 3.82E-14 | 32/157 20.3% | 537/14881 3.6% |
| GO:0005773 | CC | vacuole | 8.06E-03 | 25/157 15.9% | 1167/14881 7.8% |
| GO:0005886 | CC | plasma membrane | 1.96E-02 | 52/157 33.1% | 3358/14881 22.5% |
| **‘overstimulated’ vs. ‘control’** |  |  |  |  |  |
| GO:0019748 | BP | secondary metabolic process | 5.51E-13 | 114/1140 10.0% | 702/14881 4.7% |
| GO:0009628 | BP | response to abiotic stimulus | 1.48E-09 | 254/1140 22.2% | 2265/14881 15.2% |
| GO:0006629 | BP | lipid metabolic process | 3.41E-08 | 122/1140 10.7% | 923/14881 6.2% |
| GO:0009835 | BP | ripening | 1.02E-07 | 9/1140 0.7% | 11/14881 0.0% |
| GO:0009607 | BP | response to biotic stimulus | 2.98E-06 | 142/1140 12.4% | 1217/14881 8.1% |
| GO:0006950 | BP | response to stress | 9.35E-06 | 318/1140 27.8% | 3270/14881 21.9% |
| GO:0009605 | BP | response to external stimulus | 2.43E-05 | 186/1140 16.3% | 1760/14881 11.8% |
| GO:0005975 | BP | carbohydrate metabolic process | 1.42E-03 | 97/1140 8.5% | 879/14881 5.9% |
| GO:0009719 | BP | response to endogenous stimulus | 1.42E-03 | 188/1140 16.4% | 1918/14881 12.8% |
| GO:0006519 | BP | cellular amino acid and derivative metabolic process | 8.49E-03 | 78/1140 6.8% | 723/14881 4.8% |
| GO:0005215 | MF | transporter activity | 1.17E-04 | 133/1140 11.6% | 1210/14881 8.1% |
| GO:0016740 | MF | transferase activity | 1.80E-03 | 271/1140 23.7% | 2924/14881 19.6% |
| GO:0003824 | MF | catalytic activity | 8.48E-03 | 581/1140 50.9% | 6927/14881 46.5% |
| GO:0003674 | MF | molecular_function | 8.49E-03 | 897/1140 78.6% | 11153/14881 74.9% |
| **‘overstimulated’ vs. ‘stimulated’** |  |  |  |  |  |
| GO:0019748 | BP | secondary metabolic process | 6.07E-18 | 137/1303 10.5% | 702/14881 4.7% |
| GO:0006519 | BP | cellular amino acid and derivative metabolic process | 2.86E-11 | 122/1303 9.3% | 723/14881 4.8% |
| GO:0006629 | BP | lipid metabolic process | 2.23E-07 | 132/1303 10.1% | 923/14881 6.2% |
| GO:0009835 | BP | ripening | 3.41E-07 | 9/1303 0.6% | 11/14881 0.0% |
| GO:0005975 | BP | carbohydrate metabolic process | 2.86E-06 | 122/1303 9.3% | 879/14881 5.9% |
| GO:0009628 | BP | response to abiotic stimulus | 3.90E-06 | 263/1303 20.1% | 2265/14881 15.2% |
| GO:0009058 | BP | biosynthetic process | 6.48E-04 | 281/1303 21.5% | 2609/14881 17.5% |
| GO:0009607 | BP | response to biotic stimulus | 2.92E-03 | 141/1303 10.8% | 1217/14881 8.1% |
| GO:0009605 | BP | response to external stimulus | 1.46E-02 | 188/1303 14.4% | 1760/14881 11.8% |
| GO:0030312 | CC | external encapsulating structure | 1.46E-02 | 67/1303 5.1% | 537/14881 3.6% |
| GO:0005618 | CC | cell wall | 1.46E-02 | 67/1303 5.1% | 537/14881 3.6% |
| GO:0005576 | CC | extracellular region | 3.86E-02 | 61/1303 4.6% | 504/14881 3.3% |
| GO:0003824 | MF | catalytic activity | 3.66E-06 | 694/1303 53.2% | 6927/14881 46.5% |
| GO:0016740 | MF | transferase activity | 6.98E-03 | 301/1303 23.1% | 2924/14881 19.6% |
| GO:0005215 | MF | transporter activity | 2.92E-02 | 132/1303 10.1% | 1210/14881 8.1% |
| GO:0003674 | MF | molecular_function | 4.70E-02 | 1013/1303 77.7% | 11153/14881 74.9% |

In ‘stimulated’ *vs.* ‘control’ comparison, the main enriched GO categories are “cell wall” (GO:0005618) and “vacuole” (GO:0005773) for “Cellular Component” ontology, and “secondary metabolic process” (GO:0019748) and “cellular amino acid and derivative metabolic process” (GO:0006519) for “Biological Process”.

On the other hand, the DETs found in the two comparisons involving the multi-dropping samples ‘overstimulated’ *vs.* ‘control’ and ‘overstimulated’ *vs.* ‘stimulated’ share most of the enriched GO categories. In both genes subsets, “transferase activity” (GO:0016740) and “transporter activity” (GO:0005215) are the over-represented subcategories for “Molecular Function” ontology, while, for “Biological Process” ontology, they share “secondary metabolic process” (GO:0019748), “cellular amino acid and derivative metabolic process” (GO:0006519), “lipid metabolic process” (GO:0006629), “ripening” (GO:0009835), “carbohydrate metabolic process” (GO:0005975), “response to abiotic stimulus” (GO:0009628), “response to biotic stimulus” (GO:0009607) and “response to external stimulus” (GO:0009605) as enriched categories. The most significantly enriched “Biological Process” GO categories (adjusted *p*-value < 10^-7^) in the two comparisons are “lipid metabolic process” (GO:0006629), “secondary metabolic process” (GO:0019748), “ripening” (GO:0009835) and “response to abiotic stimulus” (GO:0009628). Lipid metabolism plays a crucial role in plant stress responses taking part to several defense process (Bhattacharya 2022), modulating membrane composition, producing stress-related hormones (i.e. abscisic acid), storing energy reserves, and protecting against oxidative damage. In particular, the notable transcripts linked to lipid metabolic process and highlighted in both the comparisons involving the multi-dropping samples were TRINITY_DN3421_C0_G1_I1, TRINITY_DN3421_C0_G1_I2, TRINITY_DN3421_C0_G1_I3, annotated as Alpha-dioxygenase 2, which is described to be involved in the oxidative degradation of fatty acids and in the formation of oxylipins (Blée 2002), implicated in signaling and stress response; TRINITY_DN2840_C0_G1_I1, TRINITY_DN2840_C0_G1_I2, annotated as the protein 3-like responsible to the elongation of fatty acids; TRINITY_DN414_C0_G1_I3, annotated as Lysophospholipid acyltransferase LPEAT2 (Jasieniecka-Gazarkiewicz et al. 2016), a protein involved in the restructuring of phospholipids and in the regulation of the fluidity and functionality of cell membranes; TRINITY_DN4194_C0_G1_I1, annotated as a chloroplastic isoform of omega-6 fatty acid desaturase, which catalyzes the introduction of double bonds into fatty acids, promoting the synthesis of polyunsaturated fatty acids essential for membrane structure and lipid signaling; and finally, TRINITY_DN455_C0_G2_I1, annotated as 3-ketoacyl-CoA synthase 1 (Luo et al. 2024), a key enzyme in the biosynthesis of very long chain fatty acids, important for cuticular waxes and structural lipids.

“Biological Process” GO categories “response to stress” (GO:0006950) and “response to endogenous stimulus” (GO:0009719) are enriched specifically in ‘overstimulated’ *vs.* ‘control’ DETs, while “biosynthetic process” (GO:0009058) is specifically enriched in ‘overstimulated’ *vs.* ‘stimulated’ comparison. This DETs subset also presents over-represented “external encapsulating structure” (GO:0030312) and “cell wall” (GO:0005618) “Cellular component” GO categories.

Regarding the KEGG pathway enrichment analysis, the number of enriched KEGG pathways was 6, 17 and 30 for ‘stimulated’ *vs.* ‘control’, ‘overstimulated’ *vs.* ‘control’ and ‘overstimulated’ *vs.* ‘stimulated’ pairwise comparisons, respectively (Table S5), and the top 10 enriched categories for each comparison were shown in Fig.3.

**Figure 3.**
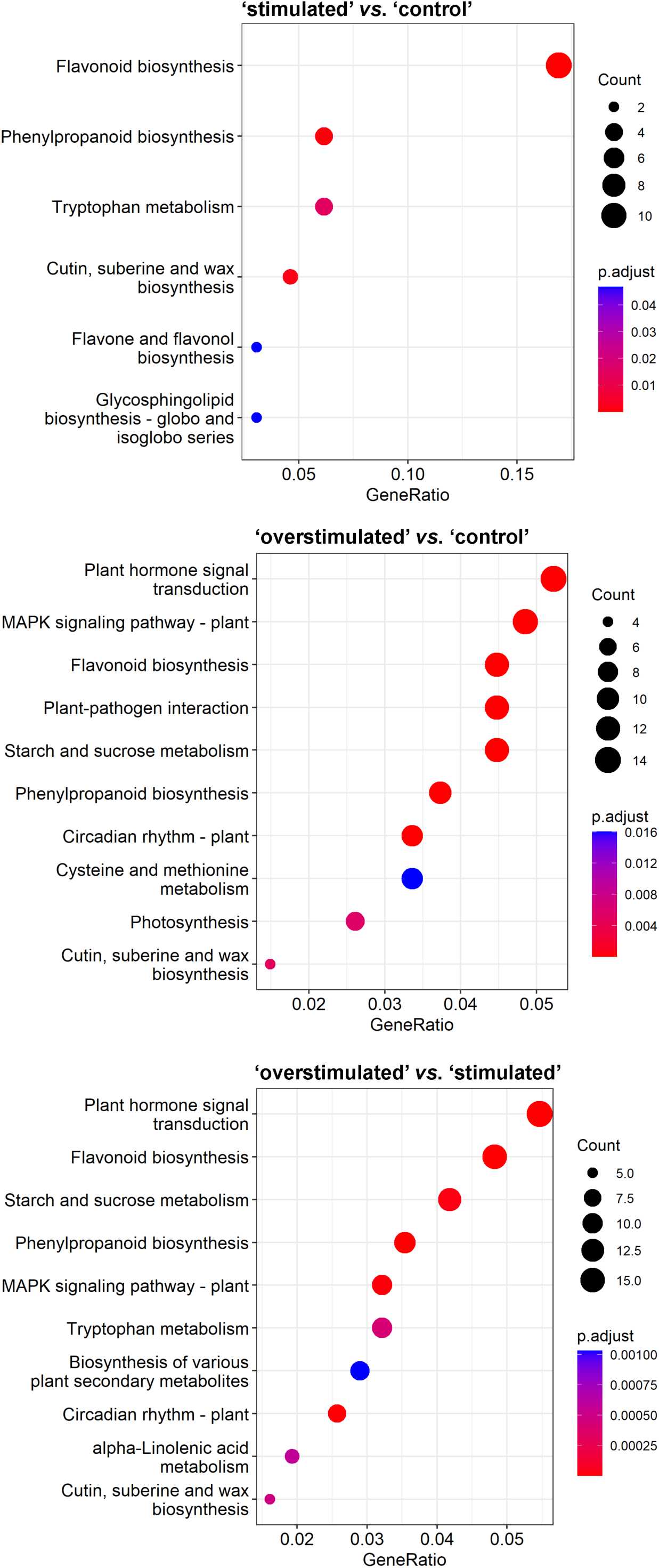
Top ten enriched KEGG pathways for ‘stimulated’ *vs.* ‘control’, ‘overstimulated’ *vs.* ‘control’ and ‘overstimulated’ *vs.* ‘stimulated’ pairwise comparisons. X axis represent the enrichment ratio (number of DETs belonging to the KEGG category / number of genes belonging to the same KEGG category in the background genome); Y axis represent the enriched KEGG categories ordered by the number of detected DEGs; the dots size represents the numbers of DETs included in each KEGG category; the dots color represents the adjusted p-value for each KEGG category enrichment.

As evidenced, there were fewer enriched KEGG pathways with the identified DETs when comparing ‘stimulated’ *vs.* ‘control’ than when comparing ‘overstimulated’ *vs.* ‘control’ and ‘overstimulated’ *vs.* ‘stimulated’. Interestingly, the “flavonoid biosynthesis pathway” category is significantly enriched, reaching a GeneRatio > 0.005 (number of DETs belonging to the KEGG category / number of genes belonging to the same KEGG category in the background genome). Contrarily, the DETs found in ‘overstimulated’ *vs.* ‘control’ and ‘overstimulated’ *vs.* ‘stimulated’ comparisons originated a greater number of enriched KEGG pathways, most of which are shared between the two subsets. In particular, “plant hormone signal transduction” (map04075), “polypropanoid biosynthesis” (map00940) and “circadian rhythm” (map04712) are among the most significantly enriched KEGG metabolic pathways in both the comparisons.

Notably, the KEGG “flavonoid metabolism” (map00941) category, which includes phenylpropanoid, flavone and flavonol, isoflavonoid and anthocyanin biosynthetic pathways, was enriched in all three comparisons, evidencing the fundamental role of flavonoids in plant response to both single or multiple stress.

Furthermore, the “plant hormone signal transduction” (map04075) category was significantly enriched (adjusted *p*-value < 0.05) in both ‘overstimulated’ *vs.* ‘control’ and ‘overstimulated’ *vs.* ‘stimulated’ comparisons, highlighting the role of multiple stress on hormones metabolic networks. In particular, map04075 includes the biosynthesis pathways of zeatin, which contribute to cytokinin biosynthesis; diterpenoid pathways, which contribute to gibberellin biosynthesis; and cysteine and methionine biosynthesis, which contribute to the ethylene biosynthetic pathway. It also includes brassinosteroid biosynthesis, jasmonic acid biosynthesis, and tryptophan metabolism, which contribute to the biosynthesis pathways of auxin and abscisic acid.

#### Characterization of genes with different transcriptional patterns

The expression level of several genes showed a similar trend in the comparisons involving plants stimulated multiple times (‘overstimulated’ *vs.* ‘stimulated’ and ‘overstimulated’ *vs.* ‘control’), and this trend was different from that observed when comparing ‘stimulated’ *vs.* ‘control’. 1,405 out of 6,714 DETs showed an up regulation in ‘overstimulated’ *vs.* ‘stimulated’ and ‘overstimulated’ *vs.* ‘control’ comparisons while they were not regulated in the ‘stimulated’ *vs.* ‘control’ comparison. Among these, TRINITY_DN1556_c0_g1_i1 (logFC = 0,28; 7,22; 6,95, for ‘stimulated’ *vs.* ‘control’, ‘overstimulated’ *vs.* ‘control’ and ‘overstimulated’ *vs.* ‘stimulated’, respectively) and TRINITY_DN1556_c0_g1_i5 (logFC = −0,25; 7,15; 7,40 for ‘stimulated’ *vs.* ‘control’, ‘overstimulated’ *vs.* ‘control’ and ‘overstimulated’ *vs.* ‘stimulated’, respectively), both showing a high divergence in expression regulation, are included in GO categories “response to stress”, “response to abiotic stimulus”, “carbohydrate metabolic process” and “biosynthetic process“. The latter category was significantly enriched only in the ‘overstimulated’ *vs.* ‘stimulated’ comparison (Table S4). In addition, an evident difference in gene expression among ‘stimulated’ *vs.* ‘control’ with respect to ‘overstimulated’ *vs.* ‘stimulated’ and ‘overstimulated’ *vs.* ‘control’ comparisons was observed for TRINITY_DN21409_c1_g2_i1, (logFC = −1,13; 4,70; 5,84; for ‘stimulated’ *vs.* ‘control’, ‘overstimulated’ *vs.* ‘control’ and ‘overstimulated’ *vs.* ‘stimulated’, respectively) included in “response to abiotic stimulus” and “response to stress”. Only 5 DETs were characterized by a down-regulation in ‘stimulated’ *vs.* ‘control’ comparison and an up-regulation in the other two comparisons. In 157 DETs, an opposite trend was observed, with an up-regulation of gene expression in the ‘stimulated’ *vs.* ‘control’ comparison and a down-regulation in the other two comparisons. Interestingly, some of these genes belong to “phenylpropanoid biosynthesis”, “flavonoids biosynthesis” (Fig. 4a). Some of the DETs involved in these pathways are often assigned as chalcone-flavanone isomerase family protein (as TRINITY_DN2436_c0_g1_i2, TRINITY_DN2436_c0_g1_i4 and TRINITY_DN2436_c0_g1_i7).

**Figure 4.**
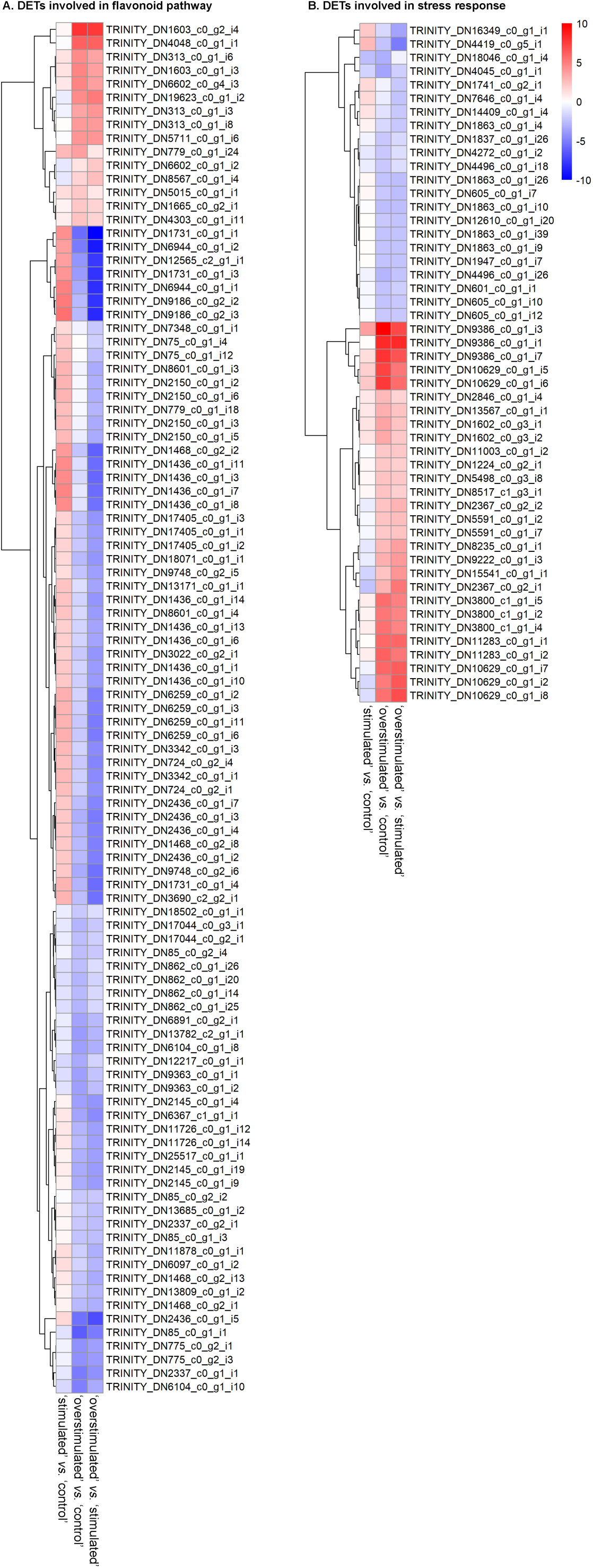
Heatmaps visualizing the log_2_ Fold Change values of the DETs involved in **A.** flavonoids pathways and **B.** biotic and abiotic stress response for ‘stimulated’ *vs.* ‘control’, ‘overstimulated’ *vs.* ‘control’ and ‘overstimulated’ *vs.* ‘stimulated’ pairwise comparisons; color key was displayed.

Among the up-regulated genes in ‘stimulated’ *vs.* ‘control’ comparison and down-regulated in the other two comparisons (‘overstimulated’ *vs.* ‘control’ and ‘overstimulated’ *vs.* ‘stimulated’) involved in “flavonoids biosynthesis”, there are TRINITY_DN11871_c0_g1_i1, TRINITY_DN12460_c0_g1_i1, TRINITY_DN12460_c0_g1_i2., TRINITY_DN12716_c0_g2_i1, TRINITY_DN13047_c0_g1_i1, TRINITY_DN14004_c0_g1_i1, TRINITY_DN17981_c1_g1_i4, TRINITY_DN17981_c1_g1_i5, TRINITY_DN21242_c0_g1_i1 TRINITY_DN2154_c0_g1_i1 TRINITY_DN2285_c0_g1_i2 and TRINITY_DN8071_c0_g1_i8. Interestingly, four of these (TRINITY_DN10306_c0_g1_i1, TRINITY_DN10306_c0_g1_i3, TRINITY_DN10306_c0_g1_i2, TRINITY_DN10306_c0_g1_i5) were annotated as leucoanthocyanidin dioxygenase, involved in proanthocyanins, class of polyphenols with various pharmacological properties, biosynthesis.

In the observed cases, the logFC in the ‘stimulated’ *vs.* ‘control’ comparison is completely in opposite to the logFC in the ‘overstimulated’ *vs.* ‘stimulated’ comparison, indicating that the genes respond significantly differently to the intensity and frequency of the stimulation.

There are no genes down-regulated in both ‘stimulated’ *vs.* ‘control’ and ‘overstimulated’ *vs.* ‘control’ comparisons and up-regulated in ‘overstimulated’ *vs.* ‘stimulated’, but 441 transcripts are up-regulated only in the comparison ‘overstimulated’ *vs.* ‘stimulated’ and not regulated in the other comparisons, and mostly involved in KEGG pathways referred to “plant hormone signals transduction” (map04075) (i.e. TRINITY_DN48442_c0_g1_i1, TRINITY_DN7301_c0_g1_i3, TRINITY_DN7339_c2_g1_i1), “phenylpropanoid biosynthesis” (map00940) (i.e. TRINITY_DN4232_c0_g1_i6).

Interestingly, 32 genes resulted down-regulated in all the three comparisons, indicating a broad effect of mechanical stimulation on genes expression. This pattern was observed in both multiple or single stress conditions compared to control plants, as well as in multiple *vs.* single stimulated plants. Most were identified as hypothetical or unidentified protein, but it seems that the diffused down-regulation in all the three comparison is frequent in transcriptional regulators (EXS and WRKY family), enzymes and disease resistance genes. Conversely, only 28 genes are up-regulated in all the comparisons, explaining in the same manner the effect of mechanical perturbation on unstressed or less-stressed plants. In this case, the involved genes determined the overexpression of DNA binding proteins as zinc fingers and helix-loop-helix, stress-induced proteins, transferases, and iron transporters.

In general, the most differentially regulated gene categories in the three comparisons are: 1) among transcription factors, NACs, WRKYs, UNE12-like, MYB108, MYB6 bZIPs, RAX2, MTERF15, MTERF2, ERF113-like, AP2/ERF, heat shock proteins, MYB6 which are usually up-regulated in ‘overstimulated’ *vs.* ‘control’ and ‘overstimulated’ *vs.* ‘stimulated’ comparisons. They aren’t subjected to variation in ‘stimulated’ *vs.* ‘control’ comparison; 2) several flavonoid biosynthesis-related genes, such as flavone synthase of cytochrome P450 family, dihydroflavonol-4-reductase, 2-hydroxyisoflavanone dehydratase, anthocyanidin reductase and flavanone 3-hydroxylase, were mainly up-regulated in ‘stimulated’ *vs.* ‘control’ comparison and down or not differentially regulated in the comparisons involving multi dropped plants. (Fig.4a); 3) among stress related genes related to plant’ stress response, both biotic and abiotic, universal stress-related proteins, early responsive to dehydration 15 proteins, NST1-like proteins, heat stress and heat shock proteins were found to be mostly regulated in the comparisons which include the multiple stimulation to plants and didn’t show consistent variation in ‘stimulated’ *vs.* ‘control’ comparison (Fig. 4b).

## Discussion

*Mimosa pudica* is a plant known for its capability of folding leaves in response to biotic contacts and abiotic disturbances (Bakshi et al. 2023). In particular, thigmonasty is a type of plant movement in which rapid leaf or leaflet closure occurs in response to physical touch or mechanical stimuli, as observed in carnivorous plant as *Dionaea muscipula* after insect stimulation (Braam 2005), and in the same *M. pudica* (Volkov et al. 2010a; Monshausen and Haswell 2013). The molecular and cellular electrical mechanisms that regulate leaf movements are the subject of recent studies (Stolarz and Trębacz 2021). Nevertheless, the genetic basis of these regulations is poorly understood, in particular when discriminating type, intensity, and frequency of the perceived stimulation (Volkov et al. 2010b; Hagihara and Toyota 2020). Furthermore, while several genes involved in multiple stress responses caused by different stressor conditions (salt, drought, temperature) have been largely investigated (Shao et al. 2015; Tan et al. 2023), only few data on the differential activation of genes in plants subjected to single or repetitive mechanical stimulation are available (Pan et al. 2012; Sewelam et al. 2014; Sato et al. 2024). In many cases, plants respond to abiotic and biotic stresses activating defensive pathways that can ‘prepare’ plants to respond more effectively to similar future stresses, resulting in quicker, stronger, or more persistent reactions. It is unclear whether this also applies to mechanical stresses, and in particular to the mechanism involved in signal perception and transduction that lead to the leaf folding behavior of *M. pudica* (Michmizos and Hilioti 2019).

We show that the gene regulatory network determining *M. pudica* leaf response after multiple mechanical stress was different with respect to both unstressed plants and plants subjected to a single stress event. First, at phenotypic level, the plants were tested with chlorophyll fluorescence analysis, a widely used method to test the activity of photosystem II (therefore, the photochemical efficiency) under abiotic and biotic stresses. Reduction of the maximum quantum yield of PSII in dark-adapted leaves (Fv/Fm) indicates the presence of stress affecting photochemical efficiency. Whereas, the quantum yield of PSII in illuminated leaves is a proxy to of photosynthesis and photorespiration, and of photosynthetic electron transport (Maxwell and Johnson 2000). Our measurements suggest that the maximal photochemical efficiency is not impaired by either single or repeated mechanical stress, as Fv/Fm remained high in all treatments. However, the lower quantum yield of PSII in leaves of ‘stimulated’ and ‘overstimulated’ plants under illumination, compared to ‘control’ plants, indicates a significant inhibition of photosynthesis and/or photorespiration. From these data we cannot say whether the inhibition of photosynthesis was caused by any of the three classic photosynthetic limitations: diffusive, biochemical, or photochemical (Flexas et al. 2004). The latter might seem less likely, given that Fv/Fm was similar in all dark-adapted samples. Nevertheless, ATP as a source of energy could be a significant factor influencing photochemistry of illuminated plants, limiting regeneration of ribulose bisphosphate and therefore photosynthesis (Williams and Bennett 1982). ATP is crucial for sustaining various cellular functions, including leaf movement (Williams and Bennett 1982). H+ ATPase is instrumental in activating membrane hyperpolarization that causes guard cells swelling under blue light (Inoue et al. 2010). In particular, *M. pudica* leaf folding is controlled by the available energy, being slowed down or stopped under dim light or in the dark (Roblin et al. 2024). Plasmodesmata are rich of ATP and the pulvins of *M. pudica*, where leaf movement is activated, are rich of plasmodesmata. However, H+ ATPase was not found in plasmodesmata of *M. pudica* pulvins, which may indicate different regulation of ATP content in these structures (Fleurat-Lessard et al. 1995). Despite the mechanisms regulating ATP in pulvins are still investigated, ATP might remain a good candidate as a source of energy limiting *M. pudica* leaf folding after multiple mechanical stress, and, at the same time, photosynthetic electron transport. Direct measurements or accurate estimations of ATP may be needed to verify this hypothesis.

Several genes involved in responses to stress may also play a role in regulating ATP production and utilization. For instance, genes that respond to salt, drought, and temperature stresses have been shown to influence metabolic pathways, including those generating ATP, while also regulating the photosynthetic efficiency (Krasensky and Jonak 2012; Shelake et al. 2022).

Interestingly, in line with our early sampling strategy, designed to capture the immediate molecular responses of *Mimosa pudica*, our findings highlighted a large number of differential expressed transcripts (DETs) potentially involved in the early signaling events underlying the plant’s rapid movements in all the comparisons among stress conditions. A greater number of DETs was observed in the comparison between ‘overstimulated’ *vs.* ‘stimulated’ plants compared to the DETs of ‘overstimulated’ or ‘stimulated’ *vs.* ‘control’, suggesting a progressive activation of metabolic pathways under repeated mechanical disturbance.

In the ‘stimulated’ *vs.* ‘control’ comparison, only 6 up-regulated DETs had a logFC > 9, indicating genes over-expressed in the single dropping condition. In particular, 3 of these belong to the “flavonoid biosynthesis” (KEGG map00941): TRINITY_DN17981_c1_g1_i5, a protein belonging to the O-fucosyltransferase family, TRINITY_DN13047_c0_g1_i1, a bifunctional 3-dehydroquinate dehydratase/shikimate dehydrogenase-like protein involved in the biosynthesis of aromatic amino acids from the metabolism of carbohydrates, and TRINITY_DN2285_c0_g1_i3, a leucoanthocyanidin reductase. Interestingly, however, these genes were not further up-regulated in ‘overstimulated’ plants.

Flavonoids, highly enriched in differentially expressed genes, are key phenylpropanoids like anthocyanins, flavonols, and proanthocyanidins, essential for plant growth, defense, and stress responses. They enhance resilience to drought and salt stress by activating SOD and POD enzymes to scavenge ROS, while also regulating stomatal aperture via abscisic acid, impacting photosynthesis and transpiration (Wang et al. 2021). Besides these, brassinosteroids (Planas-Riverola et al. 2019), enhance oxidative stress resistance, and tryptophan-derived compounds are able to mediate stress signaling, protecting the plant ensuring its survival under adverse conditions (Corpas et al. 2021).

The flavonoid genes up-regulated after a single mechanical stress are specifically annotated as involved in the synthesis of i) coumarate ligase (TRINITY_DN2181_c0_g1_i1, TRINITY_DN2181_c0_g1_i2), which plays a key role in lignin biosynthesis and the diversification of flavonoids and phenolics essential for plant growth and defense (Wang et al. 2016a); ii) flavanone hydroxylase (TRINITY_DN6259_c0_g1_i3, TRINITY_DN6259_c0_g1_i6, TRINITY_DN3690_c2_g2_i) and iii) flavonol synthase (TRINITY_DN3342_c0_g1_i3, TRINITY_DN3342_c0_g1_i1, TRINITY_DN724_c0_g2_i4, TRINITY_DN724_c0_g2_i1), key enzymes for flavonoid biosynthetic pathway; iv) leucoanthocyanidin reductase (TRINITY_DN2285_c0_g1_i2, TRINITY_DN2285_c0_g1_i1, TRINITY_DN6944_c0_g1_i3), which catalyzes the reduction of leucoanthocyanins into anthocyanins and other flavonoids, crucial for plant coloration and defense; v) chalcone synthase (TRINITY_DN3023_c0_g2_i1, TRINITY_DN1019_c0_g1_i2, TRINITY_DN2436_c0_g1_i7, TRINITY_DN13661_c0_g1_i1), catalyzes the first step in flavonoid biosynthesis, producing chalcones, essential intermediates for various flavonoids. Phenylpropanoids are rapidly regulated under stress, with polyphenols peaking during the hottest hours, while simpler antioxidants are produced at other times of the day (Tattini et al. 2015). Our results indicate that flavonoid biosynthesis is triggered early by a single stress but not enhanced by repeated stress. Under multiple stresses, plants may shift energy from costly defenses like flavonoids to other survival pathways (Simms and J. Triplet 1994; Pissolato et al. 2024).. In our specific case, ‘overstimulated’ plants may have shifted carbon and energy from flavonoid production to other pathways critical for long-term survival. Consistently, the genes involved in response to stress (biotic, abiotic, and external stimuli, according to the enriched GO categories GO:0009628, GO:0009607 and GO:0009605) were mainly up-regulated in ‘overstimulated’ plants.

Accordingly, repeated stress can lead to desensitization of the classical plant’s signaling systems, reducing activation of the genes involved in flavonoid pathway, while determining the up-regulation of several transcription factor (TF), such as WRKYs, MYB, AP2/ERF and heat shock TFs (Meraj et al. 2020). Repeated stress may enhance TF activation, enabling coordinated gene regulation (Joshi et al. 2016; Ma et al. 2022) that strengthens defenses, supports adaptation, and increases response plasticity.

Further inspection of the ‘overstimulated’ *vs.* ‘stimulated’ results showed the up-regulation of 441 transcripts that are not responding to the single stress event. These genes are mostly involved in hormone signals transduction, and some of these also in the phenylpropanoid biosynthesis, confirming the role of these molecules in mediating stress responses (Takahashi et al. 2019). This result may indeed indicate that hormonal activation may occur as a second step in the sequence, leading to the adaptation to stress (absence of leaf movement, in the case of *Mimosa*).

Some up-regulated genes may be crucial for memory-related processes involved in the response to repetitive stress. Recently, the role of transcriptional memory in response to acclimation to abiotic or biotic stress was explored, focusing on the differences among reception of a single stimulus (Type I) or of recurrent stimuli (Type II). Dynamic changes in TFs regulation and chromatin accessibility were identified as fundamental to trigger stress mnemonic mechanism (Pratx et al. 2024). We also observed an up-regulation of genes related with “lipid metabolic processes” and “carbohydrate metabolic processes” (Fig. 3), which may indicate the activation of more intricate and structural metabolic pathways for managing long-term stress memory.

The leaf folding response of *Mimosa pudica* to mechanical disturbances highlights its significance as a model organism for studying leaf responses under mechanical stress in plants. Our investigation into the gene regulatory networks involved in stress-induced leaf closure uncovered many transcriptomic differences between single and multiple stress events. Notably, most identified differentially expressed transcripts are associated with flavonoid biosynthesis, particularly when plants were subjected to a single stress event, while pathways related to abiotic stress responses were mainly regulated as a form of long-term stress response following repetitive stimulation. These findings not only enhance our understanding of the molecular mechanisms underlying plant responses to mechanical stress but also provide valuable insights into the adaptive strategies employed by *M. pudica* in fluctuating environments. To gain a more comprehensive understanding of the mechanically-induced changes in plant behavior or physiology, the transcript levels in the pulvinus should have been measured, or should be considered in future studies. All together, these knowledge paves the way for potential genetic improvement approaches by identifying candidate genes and pathways that could be targeted to enhance long-term or multi-stress resilience in plants. Understanding the molecular mechanisms underlying these processes, opens avenues for various stress management strategies, including biotechnological interventions such as genetic modifications or selective breeding, to develop crops with improved responses to mechanical stimuli, better adaptation to environmental challenges, and enhanced survival strategies in changing ecosystems.

## Data Availability

RNA-Seq raw reads data have been deposited and made available at EMBL-EBI under the accession number E-MTAB-14230.

## Abbreviations

TF: transcription factors
PSII: photosystem II
DETs: differentially expressed transcripts
ROS: Reactive oxygen species
TCH: touch-induced

## Contributions

MB: Data curation; Formal analysis; Conceptualization; Validation; Writing – original draft; Writing – review and editing; AC: Conceptualization; Validation; Writing – original draft; Writing – review and editing; CV: Methodology; LR: Conceptualization; Methodology; SP: Methodology; Formal analysis; FL: Methodology; Formal analysis; Writing – original draft; Writing – review and editing; SM: Writing – original draft; Writing – review and editing; FM: Conceptualization; Funding acquisition; Writing – review and editing

## Consent for publication

All the authors whose names appeared on the submission approved the version to be published.

## Supporting Information

**Fig. S1.** Multidimensional scaling (MDS) plot for the eight RNA libraries. Blue, green and red triangles represent ‘control’, ‘stimulated’ and ‘overstimulated’ plants, respectively.

**Fig. S2.** Venn diagram of up- and down-regulated transcripts for the comparisons ‘stimulated’ *vs*. ‘control’ (TvsC), ‘overstimulated’ *vs*. ‘control’ (MvsC) and ‘overstimulated’ *vs*. ‘stimulated’ plants (MvsT).

**Fig. S3.** Gene ontology (GO) enrichment analysis results for DETs obtained from ‘stimulated’ *vs.* ‘control’, ‘overstimulated’ *vs.* ‘control’ and ‘overstimulated’ *vs.* ‘stimulated’ pairwise comparisons. The circles are shaded based on significance level, and their radius is proportional to the number of DETs included in the corresponding GO category. Color key for adjusted *p*-value value was displayed.

**Table S1.** DETs differential expression analysis results for ‘stimulated’ *vs.* ‘control’, ‘overstimulated’ *vs.* ‘control’ and ‘overstimulated’ *vs.* ‘stimulated’ pairwise comparisons. For each DET, the log_2_(Fold Change), the log_2_(Counts per Million), the Likelihood Ratio, the *p*-value and the False Discovery Rate values were reported.

**Table S2.** DETs functional annotation. For each subject database used with blastx-fast analysis, the best hit, its description and the e-value were reported.

**Table S3.** GO and KEGG terms assigned to the *M. pudica* assembled transcriptome. The transcripts with no assigned terms were not reported.

**Table S4.** Enriched KEGG pathways for DETs obtained from ‘stimulated’ *vs.* ‘control’, ‘overstimulated’ *vs.* ‘control’, and ‘overstimulated’ *vs.* ‘stimulated’ pairwise comparisons. Each record reports the KEGG pathway ID, its description, the ratio of genes in the pathway to the total number of genes (GeneRatio), the ratio of genes in the pathway to the total number of background genes (BgRatio), enrichment *p*-value, adjusted *p*-value, *q*-value, and the orthologues associated with the pathway.

## Supporting information

Multidimensional scaling (MDS) plot for the eight RNA libraries

DETs differential expression analysis results for pairwaise comparisons.

DETs functional annotation. For each subject database used with blastx-fast analysis, the best hit, its description and the e-value were reported.

GO and KEGG terms assigned to the M. pudica assembled transcriptome. The transcripts with no assigned terms were not reported.

Enriched KEGG pathways for DETs

Enriched KEGG pathways for DETs

Gene ontology (GO) enrichment analysis results for DETs obtained from pairwaise comparisons.

Venn diagram of up- and down-regulated transcripts for the comparisons

