## Supplementary figures and images for "The transcriptional mechanism behind *Mimosa pudica* leaf folding in response to mechanical disturbance"

### Gene ontology (GO) enrichment analysis results for DETs obtained from pairwaise comparisons.

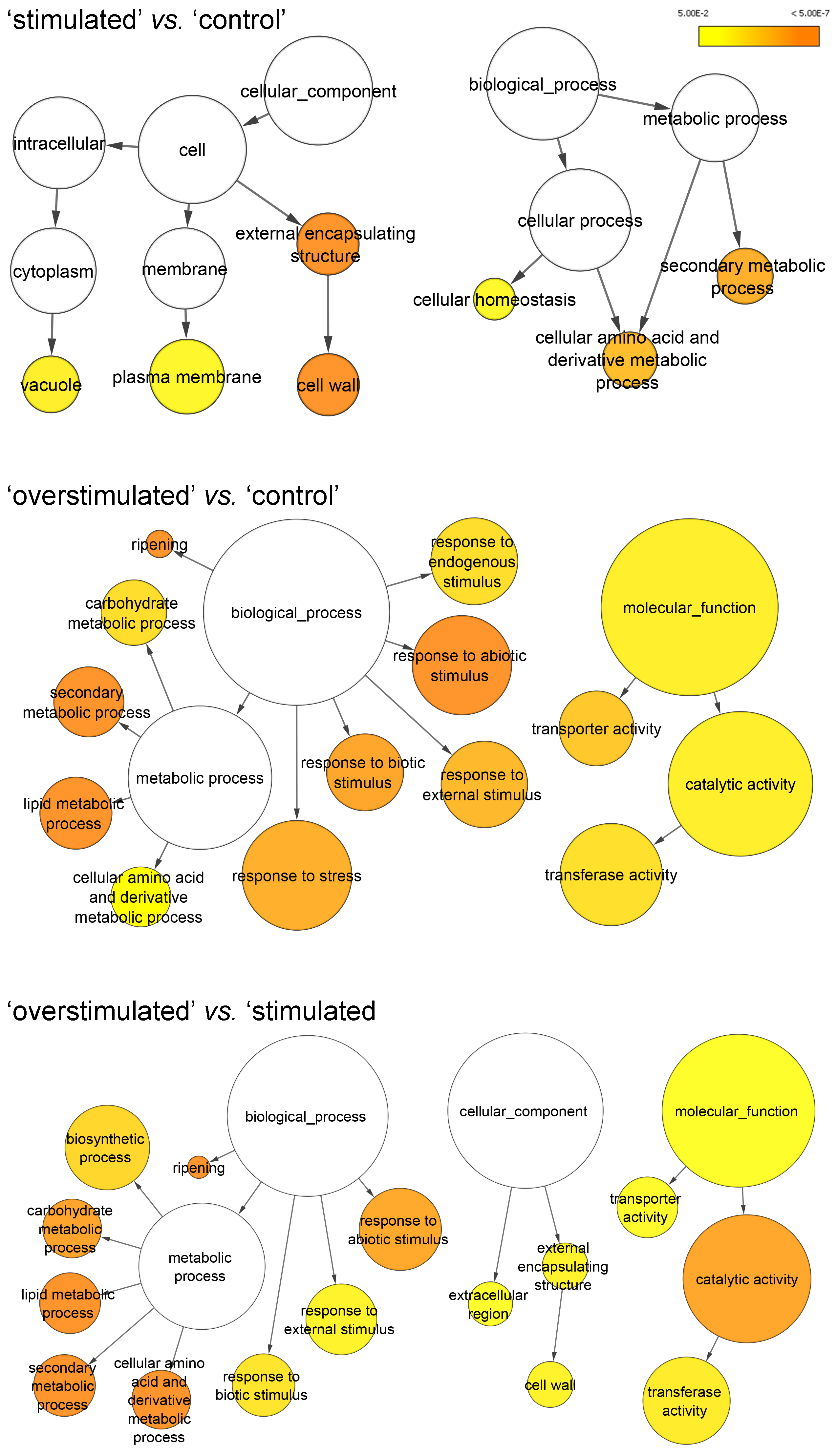

### Multidimensional scaling (MDS) plot for the eight RNA libraries

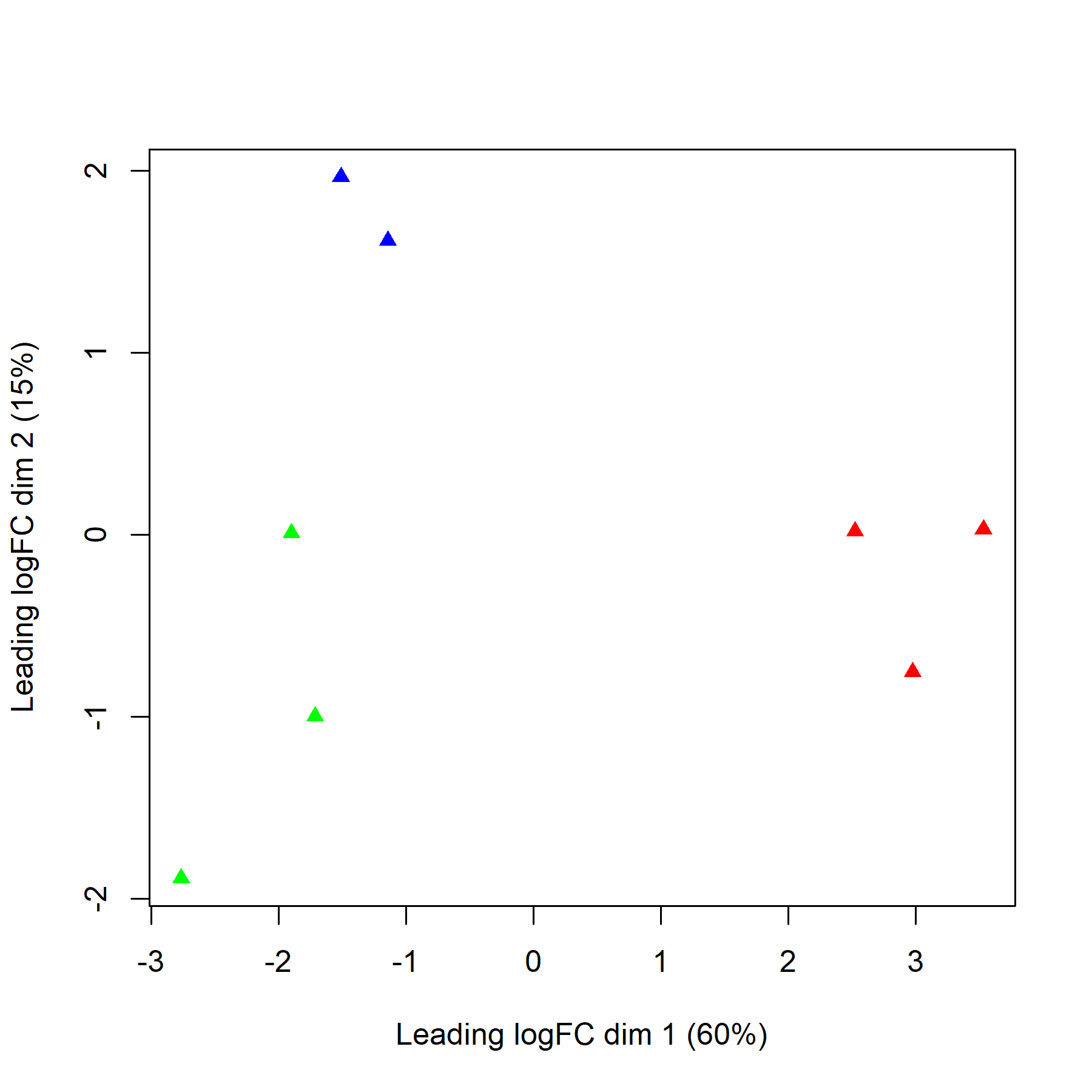

### Venn diagram of up- and down-regulated transcripts for the comparisons

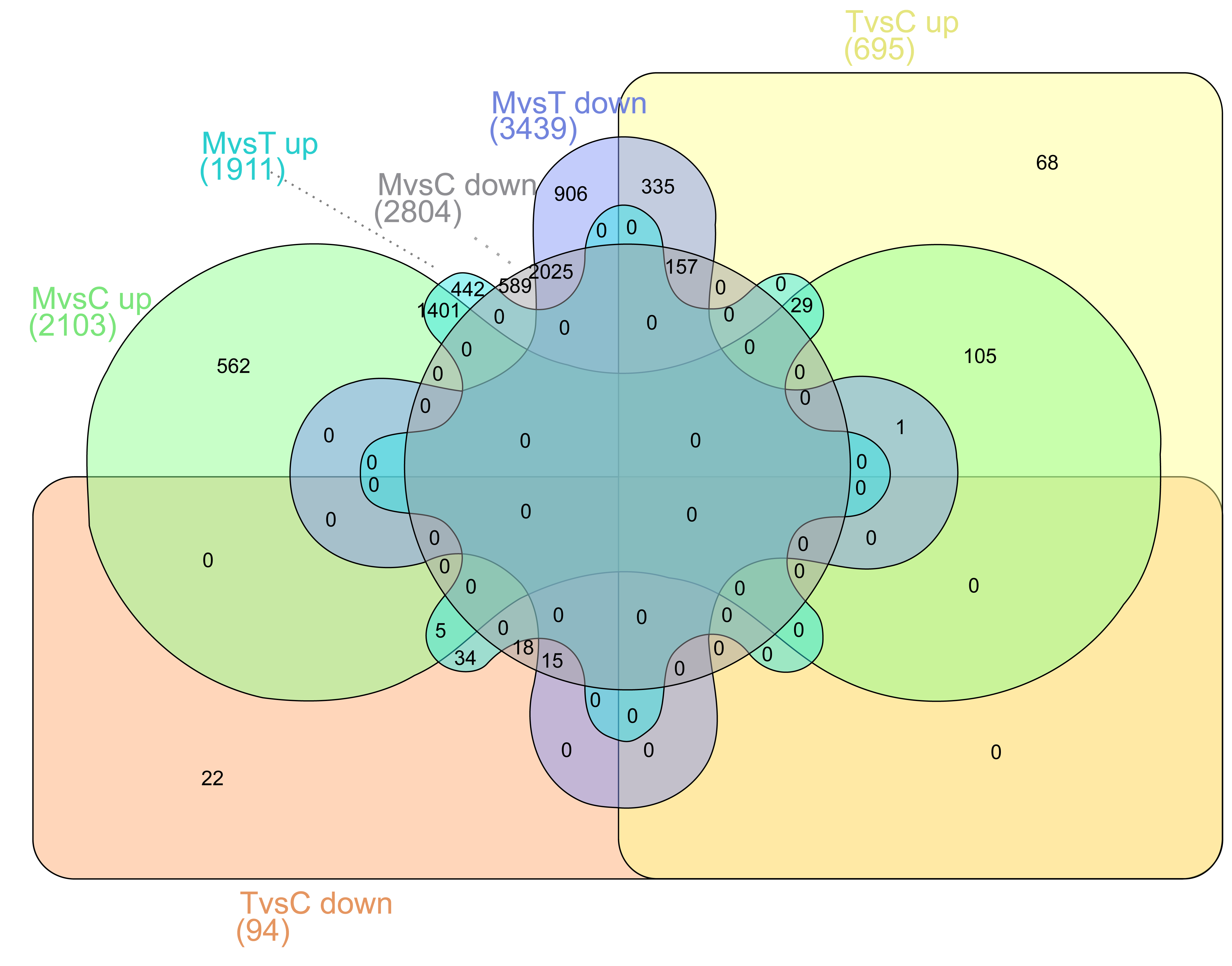
